# A fully synthetic three-dimensional human cerebrovascular model based on histological characteristics to investigate the hemodynamic fingerprint of the layer BOLD fMRI signal formation

**DOI:** 10.1101/2024.05.24.595716

**Authors:** Mario Gilberto Báez-Yáñez, Wouter Schellekens, Alex A. Bhogal, Emiel C.A. Roefs, Matthias J.P. van Osch, Jeroen C.W. Siero, Natalia Petridou

## Abstract

Recent advances in functional magnetic resonance imaging (fMRI) at ultra-high field (≥7 tesla), novel hardware, and data analysis methods have enabled detailed research on neurovascular function, such as cortical layer-specific activity, in both human and nonhuman species. A widely used fMRI technique relies on the blood oxygen level-dependent (BOLD) signal. BOLD fMRI offers insights into brain function by measuring local changes in cerebral blood volume, cerebral blood flow, and oxygen metabolism induced by increased neuronal activity. Despite its potential, interpreting BOLD fMRI data is challenging as it is only an indirect measurement of neuronal activity.

Computational modeling can help interpret BOLD data by simulating the BOLD signal formation. Current developments have focused on realistic 3D vascular models based on rodent data to understand the spatial and temporal BOLD characteristics. While such rodent-based vascular models highlight the impact of the angioarchitecture on the BOLD signal amplitude, anatomical differences between the rodent and human vasculature necessitate the development of human-specific models. Therefore, a computational framework integrating human cortical vasculature, hemodynamic changes, and biophysical properties is essential.

Here, we present a novel computational approach: a three-dimensional VAscular MOdel based on Statistics (3D VAMOS), enabling the investigation of the hemodynamic fingerprint of the BOLD signal within a model encompassing a fully synthetic human 3D cortical vasculature and hemodynamics. Our algorithm generates microvascular and macrovascular architectures based on morphological and topological features from the literature on human cortical vasculature. By simulating specific oxygen saturation states and biophysical interactions, our framework characterizes the intravascular and extravascular signal contributions across cortical depth and voxel-wise levels for gradient-echo and spin-echo readouts. Thereby, the 3D VAMOS computational framework demonstrates that using human characteristics significantly affects the BOLD fingerprint, making it an essential step in understanding the fundamental underpinnings of layer-specific fMRI experiments.

## 1. INTRODUCTION

A widely used functional magnetic resonance imaging (fMRI) technique relies on the blood oxygen level-dependent (BOLD) signal [Ogawa et al.,1993; Bandettini et al., 1994, 1997]. The BOLD signal is generated through the combined effects of changes in local deoxygenated-oxygenated blood, cerebral blood volume (CBV), cerebral blood flow (CBF) and oxygen metabolism (CMRO2) induced by neuronal activity [Ogawa et al.,1993; Bandettini et al., 1994, 1997; Belliveau et al., 1990; Uludağ et al., 2018].

Recent advances in ultra-high field MRI (≥7 tesla), novel hardware and fMRI data analysis methods, allow for the investigation of the cortical and neurovascular function at a high level of detail, e.g. at the layer-specific activity, in both human and nonhuman species [De Martino et al., 2013; Goense et al., 2006; Choi et al., 2022; Fracasso et al., 2018; Gülban et al., 2024; Huber et al., 2017; Kashyap et al., 2018; Vizioli et al., 2021; Siero et al., 2011, 2013; Bause et al., 2020; Pfaffenrot et al., 2021].

While BOLD fMRI offers significant potential for enhancing our understanding of brain function at the spatial scale of cortical layers, it is important to note that the BOLD signal is only an indirect measure of brain activity. This indirect mapping comprises a mixture of effects stemming from hemodynamic changes, the vascular architecture within the sampled volume, and the biophysical interaction between oxygen saturated blood and tissue [Uludağ et al., 2009]. Given that BOLD fMRI measures neuronal activation through hemodynamics, its ultimate spatial and temporal resolution and specificity are dictated by the spatial distribution of hemodynamic changes within the cortical angioarchitecture, and how these changes evolve over time, i.e. the hemodynamic fingerprint of the BOLD signal [Zhao et al., 2006; Siero et al., 2011].

Computational modeling of the BOLD signal formation has a long story. Starting with the simulation framework from Ogawa et al [Ogawa et al.,1993]. This model was developed to understand the susceptibility effect induced by deoxygenated blood and the macroscopic scale blobs of tissue activity, using a single cylinder model with a predefined angular orientation to mimic cerebral vessels [Ogawa et al.,1993].

More robust and complete computational frameworks succeed it using microspheres and more complex arrangements of randomly placed oriented cylinder (ROC) models within a voxel. These complex ROC models aimed to disentangle the macro- and micro-vascular influences on the BOLD signal, including the impact of pulse sequence choice on BOLD response amplitudes and vessel size specificity [Fujita N., 2001; Weisskoff et al., 1994; Boxerman et al., 1995; Yablonskiy et al., 2010; Bieri et al., 2007; Pflugfelder et al., 2011]. Moreover, computational approaches have enhanced our understanding of MRI signal characteristics. For example, they have demonstrated the relationship between biophysical interactions, such as the motion of water molecules diffusing within tissue and the susceptibility-induced effects from changes in vascular oxygen saturation levels at a mesoscopic scale [Kiselev et al., 1999, 2018; Kiselev V., 2001; Chausse et al., 2024].

Thereby, computational modeling has emerged as a significant research field aiding the understanding of BOLD fMRI signal formation, offering a means to test hypotheses in ways that experimental investigations may not always facilitate. It provides comprehensive insight into the interplay between the cerebral vasculature, intrinsic biophysical properties of the tissues, and hemodynamic changes. This is particularly relevant for ultra-high magnetic fields and high spatial resolution fMRI, e.g. submillimeter imaging resolutions, as the signal formation is more specific to certain sub-regions within the cortex that differ at the mesoscopic scale, for instance, in vascular density and architecture [Olman et al., 2012; El-Bouri et al., 2015].

Nevertheless, the impact of the three-dimensional (3D) vascular topology, associated hemodynamics, and their interaction with neighboring tissue on signal formation at the mesoscopic scale, and the temporal features of the BOLD signal evolution remain elusive [Norris et al., 2019; Norris D., 2012; Polimeni et al., 2018; Petridou et al., 2010; Dumoulin et al., 2018; Dumoulin S., 2017; Poplawsky et al., 2019; Roefs et al., 2024; Schellekens et al., 2023].

Cortical blood vessels in the human brain are organized into well-defined structures with repetitive topologies [Duvernoy et al., 1981]. These structures consist of a tree-like arrangement of penetrating arteries surrounding a central draining vein that collects deoxygenated blood from the capillary bed toward the superficial pial veins [Cassot et al., 2009, 2010; Weber et al., 2008; Schmid et al., 2019; Reichold el al., 2009; Lauwers et al., 2008; Keller et al., 2011; Hirsch et al., 2012]. In contrast to the more simplified nonrealistic vascular networks, such as ROC models, computational simulations utilizing realistic 3D vascular models extracted from the mouse parietal cortex via two-photon microscopy have highlighted the significant influence of the vascular architecture and vessel orientation on the measured BOLD signal amplitude with respect to the main magnetic field [Gagnon et al., 2015; Báez-Yánez et al., 2017, 2023]. These findings have also been corroborated by experimental data [Viessmann et al., 2019; Fracasso et al., 2018], demonstrating a phenomenon that could not be observed using nonrealistic vascular models, i.e. ROC models.

Nonetheless, realistic vascular models based on rodents [Blinder et al., 2010, 2013; Gould et al., 2017; Tsai et al., 2009] might not faithfully represent the human cortical vasculature due to interspecies differences in vascular architecture-particularly the artery/vein ratio that feeds and drains the blood in specific volumetric regions [Duvernoy et al., 1981; Schmid et al., 2019; Uludağ et al., 2018; Uludağ K., 2023] and in cortical thickness which is larger in humans than in rodents. Further, the distinct vascular densities and architectures in different cortical regions could introduce quantitative discrepancies in the simulated BOLD signals for the human brain, and potentially leading to data misinterpretation [Han et al., 2022; Lorthois et al., 2011].

In order to attain a wider understanding of the spatial and temporal hemodynamic fingerprint of the BOLD signal acquired from human brain scans, it is essential to develop a computational framework: (I) based upon the architectural layout of the human cortical vasculature, (II) that includes hemodynamic changes within this simulated vascular network, and (III) that takes the intrinsic biophysical and magnetic tissue characteristics together with MRI pulse sequence parameters into account [Gagnon et al., 2015; Markuerkiaga et al., 2021; Havlicek et al., 2017; Báez-Yánez et al., 2023; Puckett et al., 2016; Van Horen et al., 2023].

In this work, we have developed a computational framework to investigate the laminar hemodynamic BOLD signal formation based upon a fully synthetic human 3D cortical vascular model. The so-called 3D VAscular MOdel based on Statistics (VAMOS) algorithm generates both microvascular and macrovascular angioarchitectures defined by histological, morphological and topological features obtained from the human cortical vasculature [Duvernoy et al., 1981; Cassot et al., 2009, 2010; Weber et al., 2008; Schmid et al., 2019]. The microvasculature is generated through an improved Voronoi tessellation algorithm [Park H., 2021; Báez-Yánez et al., 2023] and kernel functions, while the macrovasculature is generated by kernel functions. Both vessel compartments depend on statistical properties taken from literature values, such as vessel radius, vessel tortuosity, vessel volume fraction across cortical depth, number of penetrating arteries and draining veins in a determined volume, cortical penetration dependence for large vessels, among others [Duvernoy et al., 1981; Cassot et al., 2009, 2010; Weber et al., 2008; Schmid et al., 2019]. This enables simulation of specific oxygen saturation states per vascular compartment and biophysical interactions, such as diffusion effects of water in tissue, in order to characterize the intravascular and extravascular signal contribution of diverse vascular architectures to the gradient-echo (GE) BOLD and spin-echo (SE) BOLD signals, either at the voxel level acquired at high spatial resolutions or across cortical depth, i.e. layer fMRI. The 3D VAMOS computational approach can also help to understand the impact of pulse sequence parameters on BOLD signal changes observed with submillimeter MRI acquisitions.

## 2. MATERIAL AND METHODS

### 2.1 Generation of a fully synthetic human 3D VAscular MOdel based on Statistics – 3D VAMOS model

A fully synthetic vascular model is generated using an in-house developed algorithm, and the statistical properties of the human cortical vasculature are taken from literature that estimated these by means of histology [Duvernoy et al., 1981; Cassot et al., 2009, 2010; Weber et al., 2008; Schmid et al., 2019]. First, the microvasculature (consisting of arterioles, capillaries, and venules) and the macrovasculature (comprising pial arteries and veins, penetrating arteries, and draining veins) are generated separately. Subsequently, the resulting 3D VAMOS vascular network is fully connected - by connecting the macrovascular endpoints in the arterial and venous compartments to the capillary bed, i.e. the microvascular structure. The generation process for each vascular compartment is described in the following sections.

#### 2.1.1 Generation of the microvascular architecture based on Voronoi tessellation and kernel functions

To account for the varying cortical thickness and volume fraction occupied by vessels across the human cortical grey matter, we considered that different cortical areas exhibit different cortical thicknesses and volume fractions [Fischl et al., 2000]. For instance, the primary visual cortex spans approximately 2 mm in thickness [Adams et al., 2014; Horton et al., 2018], while the primary motor cortex presents a thicker cortical depth of approximately 4 mm [Butman et al., 2007]. Therefore, initial parameters in our algorithm to be defined are a customized three-dimensional space, specifying the dimensions in x, y, and z in millimeters, and a specific vascular volume fraction, to generate the representative cortical vasculature according to the cortical region being simulated. The 3D VAMOS allows for the definition of any desired volumetric vascular dimensions, ranging from hundreds of micrometers to millimeters - thus providing versatility in creating vascular models that are not limited to represent human vascular networks but could also model mouse vascular networks, by defining the vascular properties of the studied species (see **Figure 1**). Within this volumetric space, we assumed full coverage of the cortical vasculature, extending from the superficial/pial large vessels to the cortical grey-white matter (GM-WM) boundary.

**Figure 1.**
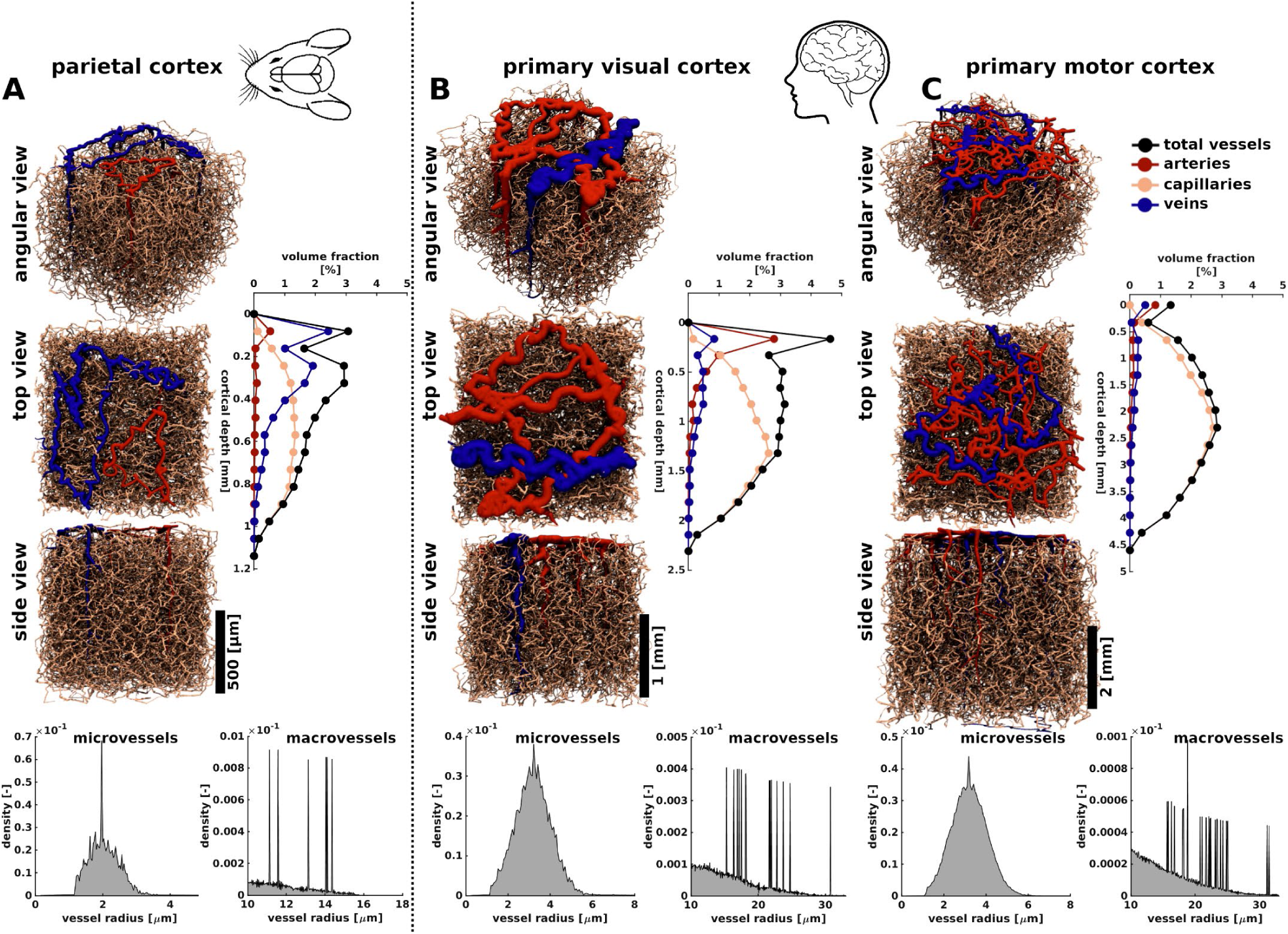
Comparison of 3D VAMOS representative mouse and human vascular models: **(A)** Representative mouse Model: We utilized the vascular characteristics described by Blinder et al. [Blinder et al., 2010] to generate a representative 3D VAMOS model of the parietal cortical vessels. Three different angular views (angular, top, and side view) are displayed, along with the respective vessel volume fraction across cortical depth. Representative human 3D VAMOS models: Taken from literature values [Adams et al., 2015; Weber et al., 2008], we present two different cortical regions, the **(B)** primary visual cortex and **(C)** primary motor cortex. The vessel volume fraction for each model are displayed along with the vessel radius distribution. Scale bar represent a 500 µm **(A)**, 1.0 mm **(B)** and 2.0 mm **(C)** cortical depth.

The microvasculature was generated using Voronoi tessellation, resulting in a topological network that resembles a mesh-like structure [Cassot et al., 2009; Lorthois et al., 2011]. Voronoi tessellations have been theoretically shown to effectively represent the capillary bed supplying brain tissue [Safaeian et al., 2011; El-Bouri et al., 2015].

The simulated volumetric space was divided into a number, S, of equidistant slabs in the xy-plane. The number S is calculated based on the vascular volume fraction and vessel features, such as vessel radius. A Voronoi tessellation was generated by fragmenting each of the slabs into tiles that encompass a given set of seed points [Park H., 2021]. The distribution of the seed points can follow any specific distribution across the slab to simulate different capillary densities across cortical depth [Schmid et al., 2019]. For example, a Gaussian distribution can be simulated in the xz-plane or yz-plane to create larger capillary densities, i.e. a larger density of Voronoi tiles in a specific part of the slab. This rational follows on that the vessel distribution across cortical depth is denser in the middle layers compared to the superficial and granular layers. [Schmid et al., 2019].

Each slab, then, was tessellated using the linear inequalities formed by perpendicular bisectors between any two connected points in the Delaunay triangulation, employing an adapted version of the polytope-bounded Voronoi diagram algorithm [Park H., 2021]. Once all the slabs contained the tessellations, joint vessels in the i-th slab were connected with the nearest (shortest Euclidean path) adjacent joint vessel of the (i+1)-th slab. This results in a fully interconnected network structure consisting of vertices, i.e. microvessel joints, and lines connecting those vertices, i.e. microvessels. Moreover, to increase complexity and mimic real capillary networks, the vertices generated by the tessellation are randomly displaced orthogonally to the slab by a small distance, typically on the order of tens of micrometers. This displacement helps create a volumetric shape for the components of each slab.

To generate a closer resemblance to actual capillary beds, an important characteristic to include is the tortuosity of the microvessels [Gould et al., 2017; Risser et al., 2007; Hartung et al., 2018]. The tortuosity (τ) was defined as the ratio of the vessel length between two joint vessel points, i.e. vertices within the Voronoi tessellation, and the Euclidean distance between those two joint vessel points (see **Figure 2**),

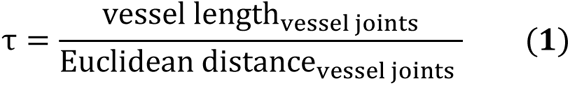

**Figure 2.**
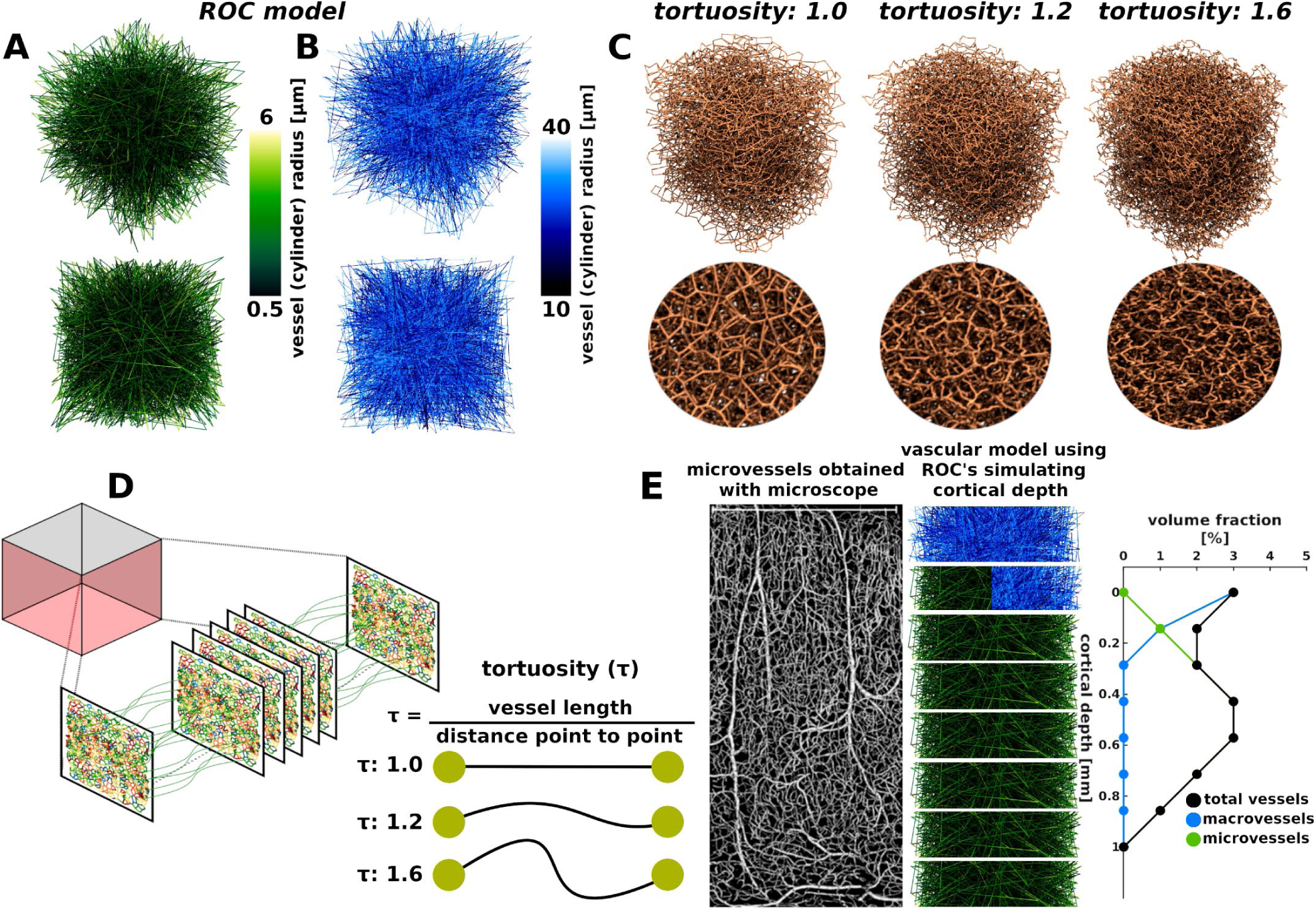
Sketch of a representative randomly oriented cylinders (ROC) model intended for visually comparing the differences in vascular architecture (**(A)**: microvessels; **(B)**: macrovessels) and a more realistic microvascular model such as the 3D VAMOS. **(C)** Examples of representative 3D VAMOS microvascular architectures for different vessel tortuosity levels. The circular images provide a zoomed-in view of each respective model. A vessel tortuosity with a value of 1.0 represents a straight line of edges as computed by the Voronoi tessellation. Tortuosity values larger than 1.0 are simulated using the kernel functions described in section 2.1.1. The model labeled as 1.6 presents a more convoluted or deformed line, resulting in a more realistic vessel topology. **(D)** Sketch of the generation of the microvascular structure as described in section 2.1.1. The slabs are tiled using a Voronoi tessellation algorithm and then connected to generate a fully interconnected network. **(E)** A schematic ROC model is shown using different vessel radius and volume fraction compositions in order to simulate cortical thickness along with a microvessel microscopy image.

In order to generate different vessel morphologies that fulfill the tortuosity characteristic, we implemented an iterative curve generator algorithm that creates plausible vessel morphologies based on predefined mathematical functions. Hereafter, we will refer to these as kernel functions. Examples of different tortuosity of the capillary bed are shown in **Figure 2**.

Along with the tortuosity, each line on the Voronoi network was assigned a value resembling the vessel radius. This value was determined by a predefined Gaussian distribution with a desired mean and standard deviation. The mean and standard deviation values were selected based on histological definitions dependent on the cortical region of investigation [Weber et al., 2008; Horton et al., 2018].

#### 2.1.2 Generation of the macrovascular architecture based on kernel functions

Based on the predefined customized three-dimensional space where the microvascular network was generated, the macrovascular architecture was constructed. The first step involves generating the pial arteries and veins. The 3D VAMOS can generate any desired number of pial arteries and veins; however, their quantity is subject to the defined number of penetrating arteries and draining veins set as initial parameters based on literature values [Duvernoy et al., 1981].

At the top plane of the volumetric space, i.e. at the maximal z-cross-section, seed points were randomly placed in the xy-plane, constrained only by the defined closer proximity value (∼120 micrometers). Subsequently, each seed point was designated to be part of an artery or a vein. The distribution of labels to the seed points follows the rationale that veins, by first principles, must be surrounded by arteries, as described by histological data from the primary visual cortex [Adams et al., 2015]. This rationale does not apply for the mouse model, given the reversed/opposite artery-vein ratio observed in mouse brain.

Next, labeled pial artery seed points were interconnected using kernel functions with predefined vessel tortuosity and vessel radius. The same process was applied to the labeled pial veins (see **Figure 3**).

**Figure 3.**
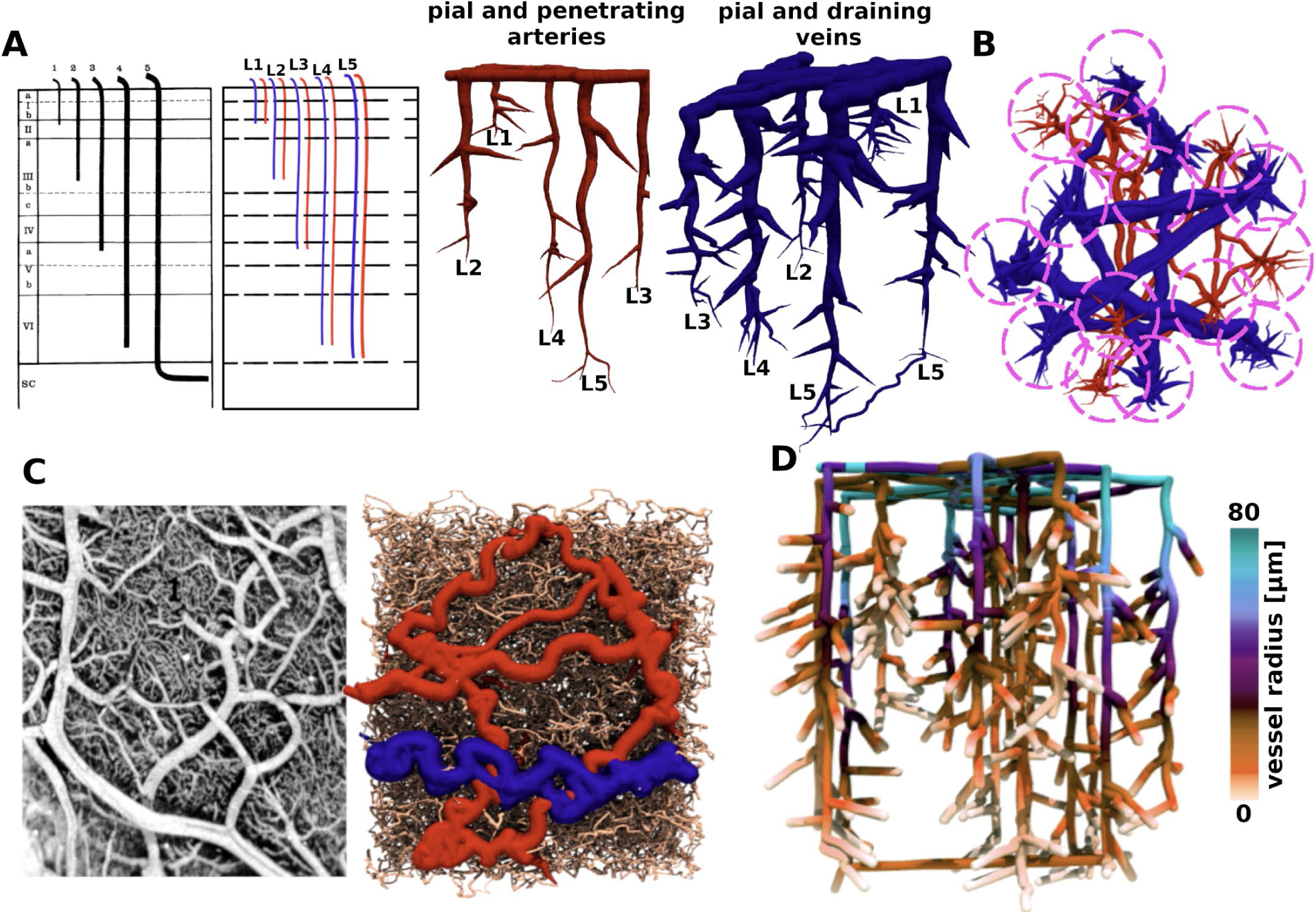
Main features accounted on the macrovascular generation. **(A)** Right: Representation of pial and penetrating arteries and pial and draining veins generated with the 3D VAMOS algorithm for different cortical penetration depths –from L1 to L5- as described by histological characteristics. The left image is adapted from [Duvernoy et al., 1981]. **(B)** Schematic representation of the radial growing of the sub-branches for each of the penetrating arteries and draining veins. **(C)** Visual comparison of a top view from the pial vasculature between the human vascular microscopy data [taken from Duvernoy et al., 1981] and the human 3D VAMOS model. **(D)** Representative macrovascular architecture showing the vessel radius distribution according to Murray’s law (R^k^parent = R^k^daughter1 + R^k^daughter2 with k = 2).

After creating the pial vasculature, the subsequent step generates the main branches of the penetrating arteries and draining veins. The 3D VAMOS facilitates the definition of the cortical penetration depth for these vessels based on the observations by Duvernoy et al [Duvernoy et al, 1981]. The main penetrating artery or draining vein can be specified to extend to varying depths, classified as Laminae 1 (L1) to Laminae 5 (L5). This classification corresponds to five equidistant laminae throughout the cortical depth (z-axis), with L1 positioned closer to the pial surface and L5 nearer to the cortical grey-white matter boundary (see **Figure 3**). Guided by the cortical penetration depth label, an endpoint of the main branch for each large vessel is aligned parallel to those of the seed points at the cortical surface and subsequently connected by another predefined kernel functions.

Furthermore, the number of sub-branches, daughters of the main vessel segment, can be predetermined as an initial parameter for each main vessel branch. These sub-branches were randomly positioned along the main vessel branch and generated using another set of predefined kernel functions. The vessel radius of both the penetrating/draining vessel and their sub-branches adheres to a branching exponent, i.e. Murray’s law (R^k^_parent_ = R^k^_daughter1_ + R^k^_daughter2_ with k values reported to be between 2 and 3 in both human and rodents; here we selected k = 2). Initially adopting the radius value assigned to the cortical pial surface seed points before gradually diminishing in radius size across the cortical depth until reaching the endpoints of the main branch and sub-branches (see **Figure 3**). Consequently, the 3D VAMOS currently generates for each parent penetrating/draining vessel (main branch) a specified number of daughter vessels expanding in a radial pattern (sub-branches), resembling a topological tree-like structure [Cassot et al., 2009]. The number of sub-branches, sub-branch vessel length and sub-branch tortuosity can be set to different values dependent on the cortical region of investigation.

One last key feature of the 3D VAMOS is the ability to define whether L5 draining veins are either connected or not at the level of the cortical grey-white matter boundary. When this parameter is selected, all labeled L5 draining veins are interconnected by predefined kernel functions (see **Figure 3**).

#### 2.1.3 Physical connection between vascular compartments

After generating both vascular compartments, i.e. the macrovessels and the microvessels, all the endpoints of the macrovascular main branches and sub-branches are connected to the nearest vessel junction of the microvascular compartment using the shortest Euclidean path between vessel joints, resulting in a fully interconnected network. This key feature allows for hemodynamic simulations, assuming only boundary conditions manipulation at the blood inlets/outlets sources, i.e. pial arteries and veins, respectively, and vasodilation changes of certain vessels or specific vascular compartments [Lorthois et al., 2011]. This improvement is significant compared to the SVM [Báez-Yáñez et al., 2023], where the macrovascular compartment was only superimposed on the microvasculature without being connected to it.

### 2.2 Simulation of the BOLD signal using the 3D VAMOS accounting for intravascular and extravascular signal contributions

The total simulated BOLD signal was calculated by summing the extravascular signal with the intravascular contributions from arteries and veins:

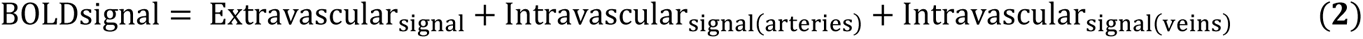

Simulations shown here were computed for gradient echo (GE) and spin echo (SE), assuming infinite readout length, at 7T using an echo time of 27 ms and 50 ms, for GE and SE, respectively, with the main magnetic field oriented parallel to the normal vector of the cortical pial surface.

#### 2.2.1 Simulation of the arterial and venous intravascular signal contribution

In this study, we assumed the intravascular contribution (R2^(∗)^_dHb_ = R2^(∗)^_0,in_ + R2_SO2_) to the BOLD signal to be non-zero for the arterial and venous compartment. This decision was based on the observation that, at high magnetic fields, the intravascular contribution of the arterial and venous compartment tends to be significant at specific oxygen saturation levels [Uludag et al, 2009]. In the microvascular compartment, we assumed zero contribution as an intravascular component, given that the R2^(∗)^_dHb_ of the capillaries has not been well-characterized due to the high heterogeneity in hematocrit levels across the cortical depth and oxygen saturation levels across the capillary bed [Gould et al., 2017].

Therefore, we implemented the intravascular intrinsic arterial and venous contribution for SE as 1/R2_0,in_ = T2_0,in_ (≈ 53 ms), and for GE as 1/R2^∗^_0,in_ = T2^∗^_0,in_ (≈ 10 ms). The R2_SO2_ component, for both pulse sequences, depends on oxygen saturation level using the quadratic relation as defined by Uludag et al. [Uludag et al, 2009], weighted by the corresponding arterial or venous blood volume fraction (see **Figure 1**).

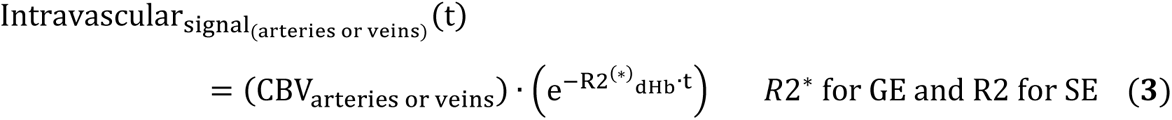

Although the intravascular decay rate for both GE and SE is influenced by hematocrit level, we assumed a constant value of hematocrit across vascular compartments in our simulations (Htc = 45%). This decision requires one degree of freedom less in the hemodynamic simulations.

#### 2.2.2 Simulation of the extravascular signal contribution

##### 2.2.2.1 Implementation of oxygen saturation levels for each vascular compartment

We simulated different oxygen saturation levels per vascular compartment, which were maintained constant over time, i.e. steady-state oxygen saturation levels were assumed. The baseline oxygen saturation (SO_2_) values used in the microvascular compartment were dependent on the oxygen saturation imposed on the veins as follows [Vovenko E., 1999]:

▄ SO_2_ in arteries (SO_2_art) = 95%;
▄ SO_2_ in capillaries = SO_2_art - ((SO_2_art - SO_2_vein) / 2);
▄ SO_2_ in veins (SO_2_vein) = [from 60% to 80% at an interval increment of 1.6%].

##### 2.2.2.2 Simulation of the extravascular signal contribution

The extravascular BOLDsignal was computed by modelling the interaction of moving spins within the local magnetic field distortions induced by the different oxygen saturation levels of both the macro- and micro-vasculature (section 2.2.2.1). We computed local frequency shifts caused by a vessel segment as the dipolar response of a finite cylinder, presuming negligible effects on the cylinder extremities [Báez-Yáñez et al., 2017]. The local frequency shift δω(r) in [1/s] for each vessel segment was computed using:

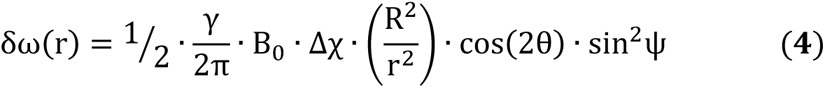

where γ is the hydrogen gyromagnetic ratio = 267.5E6 [rad/(s·T)], B_0_ is the main magnetic field (7 [T]), Δχ = 4π · 0.276 ppm · HcT · (1 − SO_2_) [-] is the susceptibility difference produced by the SO_2_in the vessel/cylinder and the hematocrit level HcT (= 0.45 [-]) [Pries et al., 1992], R is the vessel radius in [µm], r is the Euclidean distance from the center line of the cylinder to a particular spatial position in the simulation volume in [µm], θ is the angle between the cylinder and the spatial position in [rad], and ψ is the angle between the orientation of the cylinder and the main magnetic field in [rad].

The dephasing experienced by a bulk of diffusing spins (N_spins_) was simulated using a Monte Carlo approach over 20 repetitions, with 5·10^7^ spins in each repetition and assuming an isotropic diffusion coefficient of D = 1.2 [µm^2^/ms] [Kiselev V., 2001]. It is worth noting that, in order to increase the statistical averaging/power of the simulated BOLD signals, a different 3D VAMOS vascular model was generated for each repetition using the same initial vascular parameters.

Thus, the morphology of the vascular model varies in each repetition –similar to averaging the BOLD signal contribution from several voxels.

The calculation of the spin dephasing was obtained through,

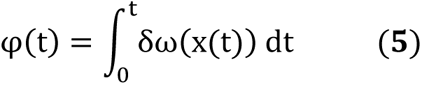

where φ(t) is the phase acquired during the simulation time t and δω(x(t)) is the local frequency shift at spin position x at each time-step t. The phasing experienced for each spin was stored across all simulation time-steps (time step = 0.025 ms). For SE sequences, the acquired phase during the echo time was multiplied by −1 (change in polarity) after TE/2, simulating the effect of the 180-degree refocusing radiofrequency pulse. Using equation **(6)** we can obtain the normalized extravascular BOLD signal.

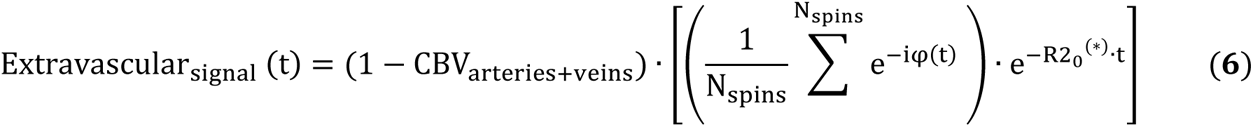

Where R2_0_^(∗)^ = 1/T2_0_^(*)^ is the intrinsic decay rate in cortical tissue, and R2’, expanded in the term inside the parenthesis, is the decay rate induced by the interaction of the diffusing spins in a local inhomogeneous frequency field. We used the intrinsic tissue T2_0_* (≈ 28 ms) relaxation time for GE and the intrinsic tissue T2_0_ (≈ 50 ms) relaxation time for SE according to the nonlinear relationship given by Khajehim et. al. [Khajehim et al., 2017] for cortical grey matter at 7T.

To confine spins within the simulation space, voxel boundary conditions were set to infinite space. Spins exiting the “voxel” re-entered the imaging volume on the opposite side, preserving their magnetization history. However, spins reaching the cortical pial surface and WM/GM boundary were considered invalid iterations and reiterations were performed. Additionally, spin exchange between vascular compartments was prohibited, establishing an impermeable vascular network.

### 2.3 GE R2* and GE BOLD signal change, and SE R2 and SE BOLD signal change across cortical depth

The BOLDsignal and the corresponding GE R2* and SE R2 were computed using their respective echo time as described in section 2.2. Given that the behavior of the MR signal, in general, presents oscillations due to its multi-exponential nature [Kiselev V., 2001], we simply approximate the R2^(^*^)^ decay rate value fitting a polynomial of degree one, i.e. a linear fit, on the natural logarithm of the BOLDsignal, 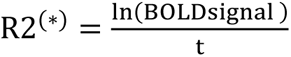, for GE and SE, respectively. The BOLD signal change in [%] was defined as the relative change using the 60% oxygen saturation state as the reference/baseline condition (SO_2vein_ = 60%), i.e.,

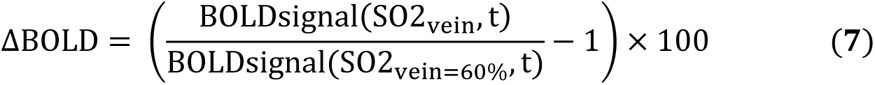

Each simulated model was divided into fifteen layers to characterize the behavior across cortical depth. These layers do not represent or resemble any realistic definition of cortical layers obtained through histological samples.

### 2.4 Using randomly oriented cylinder (ROC) models to simulate cortical layer BOLD signals

In order to demonstrate the advantages of using a realistic 3D vascular model, we generated a composite vascular model using ROCs, as displayed in **Figure 2**, to simulate the laminar BOLD contribution. Macrovascular ROC models were simulated using cylinder/vessel radius sizes ranging from 10 µm to 40 µm, and microvascular ROC models were simulated using cylinder/vessel radius sizes within the range of 0.5 µm to 6 µm, as shown in **Figure 2**. We imposed a volume fraction dependent on cortical depth for both macrovascular and microvascular ROC models. Thus, simulating a cortical thickness of 1 mm isotropic, we divided it into eight equidistant layers. The volume fraction of each layer depends on the vascular compartment. The macrovascular ROC model follows a volume fraction of 3% at the cortical surface, decreasing its value across cortical depth, as depicted in **Figure 2**. The microvascular ROC model intents to simulate a Gaussian distribution, with a 3% value in the middle layers and decreasing values toward the cortical surface and the GM/WM boundary, as shown in **Figure 2**. The oxygen saturation levels used were selected to match the values typically found in veins – SO2 = 60% to 80% (see section 2.2.2.1). Moreover, we employed the same biophysical properties of tissue as described previously.

## 3. RESULTS

To demonstrate the capabilities and versatility of the 3D VAMOS computational framework, we generated three different vascular models – one mouse vascular model (parietal cortex) and two human vascular models (primary visual cortex and primary motor cortex) (see **Figure 1**).

These 3D VAMOS models were confined to a simulation space of an approximately 1 x 1 x 1 mm³ for the mouse model (**Figure 1.A**); 2 x 2 x 2 mm³ for the primary visual cortex (**Figure 1.B**), and 4 x 4 x 4 mm³ for the primary motor cortex (**Figure 1.C**). These vascular models represent the corresponding cortical thickness of each brain region and species.

In **Figure 1.A**, three different viewing angles of the generated mouse model (angular, top, and side views) are shown. For the generation of the mouse microvascular compartment, the vessel radius was set to a mean value of 2.2 µm and a standard deviation of 0.5 µm [Blinder et al., 2013; Schmid et al., 2019]. The simulated tortuosity was set to 1.2. The vessel distribution across cortical depth followed a Gaussian distribution with a peak at the middle cortical layers (∼500 µm in depth) and gradually reduces its value towards the cortical surface and the cortical grey-white matter boundary. Upon descriptions of the macrovasculature of the mouse, the number of penetrating arteries and draining veins followed an artery-vein ratio of ∼1:3 [Blinder et al., 2013; Schmid et al., 2019]. Hence, two arteries - with radius in the range of 7 µm to 12 µm - and six veins - with radius in the range of 10 µm to 14 µm - as per 1.0 mm^2^ were used [Blinder et al., 2013; Tsai et al., 2009; Schmid et al., 2019]. All macrovessels have a cortical penetration labeled as L4, meaning that approximately all penetrating arteries and draining veins reach a penetration depth of around 70% to 85% in the model. The distribution of the vessel radius and volume fraction are depicted along the model.

Furthermore, in **Figures 1.B** and **1.C**, two different simulated human cortical regions are shown in three different viewing angles (angular, top, and side views): a representative primary visual cortex **(1.B)** and a representative primary motor cortex **(1.C)**. The models comprised a microvascular structure with a vessel radius distribution obtained by a Gaussian distribution with a mean vessel radius of 3.235 µm and a standard deviation of 0.85 µm [Weber et al., 2008; Cassot et al., 2009; Lorthois et al., 2011]. The simulated tortuosity was fixed to 1.2. The vessel distribution across cortical depth followed a Gaussian distribution with a peak at the middle cortical layers (∼1.0 mm and 2.0 mm, respectively) and slowly reduces its value towards the cortical surface and the cortical grey-white matter boundary. The human models were set to an artery-vein ratio of ∼3:1 [Duvernoy et al., 1981]. For the primary visual cortex, ten arteries -with radius in the range of 13 µm to 23 µm- and four veins -with radius in the range of 15.65 µm to 31.65 µm- as per ∼2.0 mm^2^ were set. For the primary motor cortex, twenty arteries and eight veins -with similar vessel radius as primary visual cortex- as distributed per ∼4.0 mm^2^ were implemented [Horton et al., 2018; Weber et al., 2008]. Macrovessels were set to different cortical penetrations, labeled from L3 to L5 – macrovessels labeled as L5 were connected at the cortical grey-white matter boundary and a minimum radial positioning distance between penetrating arteries and draining veins of approximately 120 micrometers for both models. The distribution of the vessel radius and volume fraction are depicted along the model.

In **Figure 2**, we present sketches of representative ROC models representing either microvascular (**Figure 2.A**; green voxel) or macrovascular (**Figure 2.B**; blue voxel) structures, allowing for a visual comparison of their morphology with the respective microvascular network generated by the 3D VAMOS algorithm. **Figure 2.C** illustrates different tortuosity level. **Figure 2.D** displays a sketch of the generation process of the 3D VAMOS microvascular compartments, as described in section 2.1.1. Additionally, to illustrate the difference between realistic 3D vascular networks and ROC models, a schematic ROC model is shown in **Figure 2.E**, simulating different vessel radius and volume fraction compositions.

In **Figure 3**, we illustrate the features and capabilities of the 3D VAMOS in generating the macrovascular architecture. **Figure 3.A** shows the different vessel cortical penetration depth for both penetrating arteries and draining veins, respectively. The vessel penetration depth follows the description reported by Duvernoy et al [Duvernoy et al., 1981]. **Figure 3.B** shows the radially growing of the sub-branches for all the macrovascular architecture. In **Figure 3.C** we qualitatively demonstrate the resemblance of the cortical pial vessel acquired with microscopic data (left image) and the 3D VAMOS model (right image). Finally, **Figure 3.D** illustrates the radius distribution, defined by the Murray’s law, for all the macrovascular compartment.

In **Figure 4**, we present a comparison of layer BOLD signal profiles across species (mouse and human) and cortical regions using distinct simulated vascular architecture characteristics, including a layer ROC model. The dotted lines of the layer-signal profiles represent the mean value and the shaded area represents the standard deviation computed through the different oxygen saturation levels computed by the Monte Carlo approach, as described in section 2.2.2.1.

**Figure 4.**
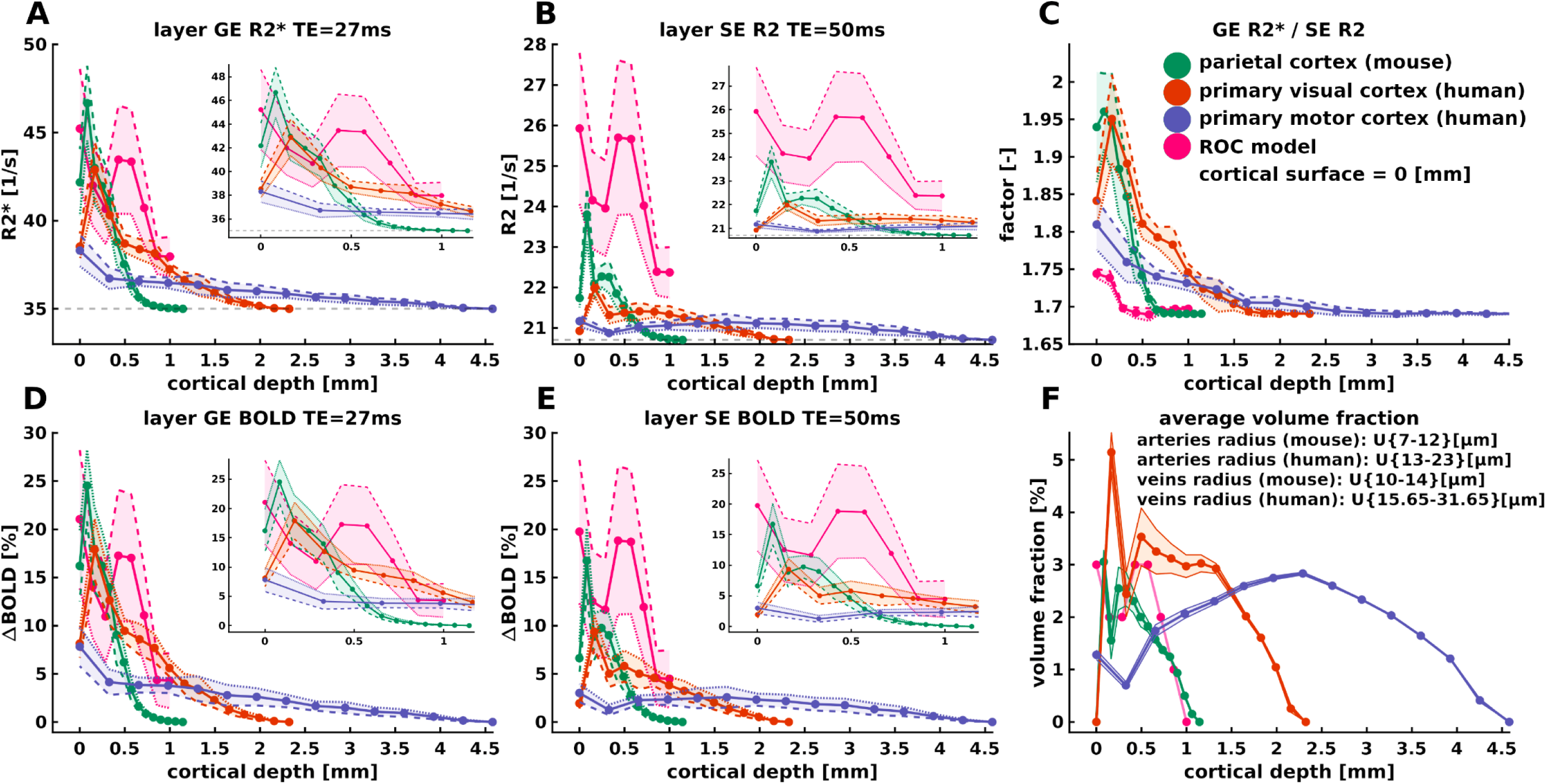
Comparison of layer BOLD signal profiles across species (mouse and human) and cortical regions using distinct simulated vascular architecture characteristics including a layer ROC model as defined in Figure 2. Dotted lines represent the mean value computed through the different Monte Carlo simulations, while the shaded area represents the standard deviation – dashed lines represent the lowest oxygen saturation value, while the segmented line represents the highest oxygen saturation value. (**A)** and (**B)** depict the R2(*) decay rate across cortical depth, respectively. **(C)** shows the GER2*/SER2 ratio as a surrogate measurement of vessel specificity. **(D)** and **(E)** present the layer BOLD signal changes across cortical depth for GE and SE, respectively; and **(F)** the average volume fraction obtained through the Monte Carlo repetitions. In mouse, the vascular architecture features a cortical depth of approximately 1.0 mm^3^, with an artery-vein ratio of 1:3 (2 arteries -with radius in the range of 7 to 12 um- and 6 veins -with radius in the range of 10 to 14 um-per 1 mm^2^), and microvessel radius parameters of mean = 2 µm and std = 0.5 µm. Human primary visual cortex simulations depict a cortical depth of around 2.0 mm^3^, with an artery-vein ratio of 3:1 (10 arteries -with radius in the range of 13 to 23 um- and 4 veins -with radius in the range of 15.65 to 31.65 um- per ∼2.0 mm^2^), and microvessel radius parameters of mean = 3.235 µm and std = 0.85 µm. Similarly, the simulated vascular architecture of the human primary motor cortex includes a cortical depth of approximately 4.0 mm^3^, an artery-vein ratio of 3:1 (20 arteries and 8 veins -with similar vessel radius as visual cortex- per ∼4.0 mm^2^), and microvessel radius parameters of mean = 3.235 µm and std = 0.85 µm. Oxygen saturation levels are described in section 2.2.2.

In **Figure 4.A**, we show the R2* [1/s] decay rate induced by each vascular model using an echo time of 27 ms. The R2* values range between ∼35-48 [1/s]. All models exhibit a larger contribution towards the cortical/superficial layer, displaying a decreasing R2* decay rate towards the GM-WM boundary –a value relatively similar to the R2* value of tissue at 7 tesla (R2* of tissue = ∼35 [1/s]). Given that the composition of the superficial vessels in the mouse model is largely comprised of veins, the mouse model shows a larger R2* towards the cortical surface compared to both human models. The ROC model displays similar R2* values compared to the 3D VAMOS models, except for the increased bump at the middle part of the model due to the larger volume fraction of microvessels used in the simulation - ROC volume fraction is displayed in **Figure 2**.

In **Figure 4.B**, the R2 [1/s] decay rate obtained by a SE readout at an echo time of 50 ms is shown. The R2 values range between ∼20-28 [1/s]. Similar behavior of layer profiles is obtained as compared to R2* decay rates. **Figure 4.C** presents the ratio of the GE R2*/SE R2 as a surrogate measurement of vessel specificity. **Figures 4.D** and **4.E** show the layer BOLD signal changes in percentage [%]. The GE BOLD signal changes range between ∼2%-30% depending on cortical depth, whereas in the SE BOLD range within an interval of ∼1% to 20% depending on cortical depth. GE BOLD signal change shows a larger contribution at the cortical surface, reducing its value across cortical depth. Similarly, SE BOLD signal changes present larger values at the cortical surface, reducing across cortical depth, except that these values are smaller compared to the GE BOLD signal changes. Finally, in **Figure 4.F**, we plot the average volume fraction that iteratively changes the morphology of each vascular model while conserving the same topological features.

## 4. DISCUSSION

### 4.1 General discussion

In order to understand the functional MRI signals obtained at high spatial resolution, i.e. at submillimeter scales, we have developed a computational framework reflecting a fully synthetic human 3D cortical vascular model obeying the MR physics governing the MR signal formation process.

The so-called 3D VAMOS algorithm generates both microvascular and macrovascular angioarchitectures defined by histological, morphological and topological features obtained from the human cortical vasculature. The microvasculature is generated through Voronoi tessellation and kernel functions, while the macrovasculature is generated by kernel functions. Both vessel compartments depend on statistical properties taken from literature values.

The computational time required to generate both macrovasculature and microvasculature depends on cortical dimensions and characteristics. Microvasculature is typically generated within seconds to a couple of minutes, while macrovasculature is generated in less than ∼2 seconds. Other computational algorithms that resemble realistic vascular architecture [Hartung et al, 2021], using a different mathematical approach, can take several days of computation, depending on vascular architecture properties and vessel characteristics. The proposed 3D VAMOS offers the advantage of relatively fast generation of representative vascular angioarchitecture. This allows for increased statistical power/ averaging of the BOLD signals, as each MR signal iteration creates a new vascular morphology while maintaining vascular topology features – similar to averaging voxels in data analysis pipelines.

Further, after generating both vascular compartments, the VAMOS algorithm results in a fully interconnected network. This crucial feature will enable an easily extension to include hemodynamic simulations with manipulation of the boundary conditions at the blood inlets/outlets sources, such as pial arteries and veins, respectively, and vasodilation changes of specific vessels or vascular compartments [Lorthois et al., 2011]. This ability is a significant improvement compared to the SVM [Báez-Yáñez et al., 2023], where the macrovascular compartment was merely superimposed on the microvasculature without being connected to it.

Nevertheless, the 3D VAMOS model enables simulation of specific oxygen saturation states per vascular compartment and biophysical interactions, such as diffusion effects of water in tissue, to characterize the intravascular and extravascular signal contribution of diverse vascular architectures to the gradient-echo (GE) BOLD and spin-echo (SE) BOLD signals, either at the voxel level acquired at high spatial resolutions or across cortical depth.

### 4.2 On the fully synthetic human 3D VAMOS model

One of the motivations for developing a fully synthetic 3D vascular model is the limitation of human cortical vasculature samples. Currently, 3D visualization of ex-vivo human vascular network samples can be achieved using immunohistochemistry labeling combined with microscopy or x-ray microtomography imaging techniques. However, these technologies are still being developed and are challenging to apply, especially for sufficiently large tissue samples. Moreover, the tissue samples may suffer from degradation and deformation. As a result, acquiring large volumes of human cortical vasculature (>1 mm^3^ isotropic) is quite difficult [Cassot et al., 2006; Lauwers et al., 2008; Duvernoy et al., 1981]. To overcome the limitations of a realistic 3D representation of the human cortical vasculature, the 3D VAMOS computational approach provides a versatile solution by generating human vascular models based on angioarchitectural characteristics derived from literature values.

Structurally, the ROC models prove inadequate in representing vascular networks when attempting to understand the formation of the BOLD signal at high spatial resolutions (see **Figure 2** and **Figure 4**). It has been demonstrated that at mesoscopic levels, the angioarchitecture adheres to well-defined patterns, such as the mesh-like network of the capillary bed. While it is indeed possible to generate ROC models by assuming monosized cylinders or a mixture of cylinder sizes within a volumetric space while maintaining the volume fraction, such vascular models cannot effectively compute specific vascular topologies, such as the well-structured penetrating arteries and draining veins [Markuerkiaga et al., 2016]. Furthermore, conducting hemodynamic simulations becomes more complex due to the high dependence of hemodynamic changes on vascular properties and topology.

Further, a detailed model of the cerebral vasculature, such as the 3D VAMOS model, is necessary to understand the underlying principles of tissue perfusion at submillimeter spatial scales. The 3D VAMOS model could be used to further our understanding of physiology. For instance, it can provide insights into the spatial distribution of oxygen by the vascular network and other hemodynamic information at any specific point within the neural tissue supplied by the vascular network [Risser et al, 2007].

Lauwers et al [Lauwers et al, 2008] observed that large vessels (macrovessels) contribute more to the vascular volume in the upper layers of the cortex, while the capillary compartment made a greater contribution in the middle third of the cortex. This cortical vascular feature can be well captured by the 3D VAMOS, as observed in the cortical depth profiles in **Figure 1**, given that the definition of the vessel decreasing vessel radius across cortical depth using the kernel functions. This distribution pattern could potentially influence the layer profiles of functional activity that have recently been observed at high resolution layer fMRI, and thus, the 3D VAMOS model is suited to investigate further the layer-specificity of this functional signal [Markuerkiaga et al., 2021; Bause et al., 2020]. Moreover, pial arteries are known to exhibit anastomoses to efficiently support regions of high perfusion demand or collateral flow [Adams et al., 2015]. However, in this manuscript, anastomoses are not implemented. Only main pial vessel segments belonging to the large feeding arteries are included. Pial veins do not exhibit anastomoses at any cortical depth level [Duvernoy et al., 1981].

It is important to note the flexibility that the 3D VAMOS offers. For example, the representative vascular models for any specific brain region are not constrained to specific macrovessel topological or morphological features. It can generate different realistic or, even, non-realistic artery-vein ratios to understand the impact of the macrovasculature on hemodynamic changes and its direct effect on BOLD fMRI signal formation.

Another advantage of the 3D VAMOS is that the vascular network is fully connected. This will allow for local hemodynamic simulations of changes in CBF, CBV, and corresponding oxygen saturation levels [Báez-Yánez et al., 2023]. This capability can help understand the specific physiological roles of the vascular compartments, their contributions to hemodynamic changes and the direct impact on the BOLD signal. For example, given its fully vascular connectivity, in future studies, we envision investigating the local transients of red blood cells in a vascular network and their effects on the heterogeneity of mean transit time [Jespersen et al., 2012], among other hemodynamic changes induced by neuronal activation or other kind of stimuli, such as controlled gas-challenges.

### 4.3 On the fully synthetic mouse 3D VAMOS model

Another motivation for developing a fully synthetic 3D vascular model is the limitation faced by advanced imaging techniques, such as two-photon microscopy or scanning electron microscopy, in capturing detailed mouse vascular structures due to the finite penetration capacity of the illumination they employ. Consequently, the depth of field of view in these methods is typically restricted to a few hundred micrometers at best. The 3D VAMOS approach helps to understand the BOLD signal formation and effects of the vascular topology and hemodynamics at the level of MRI voxels, as those acquired in fMRI measurements even at the mesoscopic laminar level.

Moreover, microscopy imaging data, such as two-photon microscopy, typically depict biological vascular structures that exhibit irregular vessel shapes. Image noise and visualization artifacts further contribute to vessel characterization degradation. Any disturbances can significantly impact the skeletonization process, often resulting in undesirable outcomes. Skeletonization is inherently sensitive to these minor boundary perturbations, necessitating the removal of unwanted effects in a post-processing stage. Distinguishing between genuine features and artifacts is often challenging, making segmentation a potential source of error in topological descriptions. The 3D VAMOS can help overcome this limitation by providing versatility in generating different vascular approximations for a wide range of vascular parameters at low computational cost and time – the generation of a fully connected vascular network can take less than ∼45 seconds, depending on the simulated vascular features and desired volumetric space. Nevertheless, refinement in post-processing microscopy data will enhance our understanding of such complex networks and provide a better-informed 3D VAMOS vascular network.

### 4.4 On the simulated BOLD fMRI signals

Given that the main vascular contribution to BOLD signal formation is attributed to the venous compartment, the mouse model exhibits larger decay rates and BOLD signal changes near the cortical surface compared to the human models –due to the artery/vein ratio. This highlights the importance of employing vascular models that replicate specific vascular features found in different species in order to reduce misinterpretations of the measured BOLD fMRI data across species.

The primary motor cortex, with an average thickness of roughly 4.0 mm in humans, presents a notable contrast to the primary visual cortex, which averages around ∼2.0 mm [Palomero-Gallagher et al, 2019]. This distinction holds significance when applying imaging and analysis methods from one regional cortex to another. Despite achieving equivalent imaging resolution in both areas, the primary motor cortex exhibits lower relative signal changes (see **Figure 4**). Hence, when comparing layer activity profiles across participants and brain regions, it is important to consider their relative cortical thickness.

For all models, GE R2* decay rates increase towards the cortical pial surface. SE R2 decay rates also increase towards the cortical pial surface, though to a lesser extent due to the refocusing 180-degree pulse. Despite this pulse, the influence of macrovessels at the cortical pial surface remains significant depending on the simulated oxygen saturation level. In the deeper layers, most contributions to GE R2* and SE R2 originate from the tissue’s R2*, with a small weighted contribution from extravascular and intravascular CBV and minor contributions from R2’.

Our findings suggest that the diverse vascular architecture in deeper gray matter has a diminished effect on the laminar signatures of both BOLD signal changes and R2* decay rates (see **Figure 4**). Conversely, superficial layers (pial surface) exhibit significant differences in the topology of large vessels, leading to a less uniform BOLD signal change in these layers. These simulation results highlight the necessity of addressing the bias of large vessels toward the pial surface in laminar fMRI data through filtering and/or normalization techniques [Vizioli et al., 2021].

### 4.5 Future studies and computational improvements

Current noninvasive functional neuroimaging methods mainly rely on detecting the hemodynamic response to neuronal activation. Improving our understanding of cortical vascular topology and functioning will enhance our insights into the effects of local cerebral blood flow disruptions on both local and global perfusions. Thus, dynamic changes of cerebral blood volume and flow will be included in future studies in order to understand the dynamic processes that drives the hemodynamic fingerprint of the BOLD signal formation and other neuroimaging techniques, such as perfusion imaging.

To enhance confidence in the resemblance of the 3D VAMOS model to realistic human vascular angioarchitecture, we intend to compare our model to various quantitative measurements in future studies. These may include distance map values of the tissue-vessel spatial distribution. Another example could be the analysis of the vascular surface-to-volume ratio.

Another methodological improvement that we will consider in the near future is the generation of vascular angioarchitecture that presents a certain degree of simulated cortical curvature. Ongoing developments include the 2D slabs, used to create the Voronoi tessellation, to be inserted in quasi-spherical spaces. The 2D slabs could be placed radially - with respect to a certain origin point to manipulate the degree of “orthogonality” with respect to the cortical surface.

In addition to this, we have assumed isotropic diffusion motion within the tissues. It has been shown that different diffusion regimes can have a strong effect on the BOLD signal, such as the one provided by the CSF [van Horen et al., 2023]. In future studies, we plan to implement a diffusion coefficient value dependent on cortical depth, i.e., CSF water motion in the superficial layers displaying a slightly different value compared to the deeper layers.

Moreover, it has been observed that penetrating arteries in certain cortical regions create cylindrical spaces devoid of capillaries in their proximity [Duvernoy et al., 1981; Cassot et al., 2010; Lauwers et al., 2008]. We plan to incorporate this realistic topological characteristic into the generation of the microvascular network in the 3D VAMOS model.

## 5. CONCLUSION

Understanding the spatial specificity of hemodynamic fingerprint BOLD fMRI signals acquired at mesoscale through a more robust and complex modeling approach, such as the one presented in the 3D VAMOS computational approach, will enhance our understanding of neuroimaging at submillimeter scales, both in healthy and pathological conditions. Therefore, the 3D VAMOS computational approach will help understand the influence of human 3D vascular architectures on the formation of hemodynamic fingerprint GE BOLD and SE BOLD signals across cortical depth and/or voxel-wise levels at high spatial imaging resolutions, as well as the impact of pulse sequence parameters on BOLD signal changes in submillimeter MRI acquisitions.

## ACKNOWLEDGEMENTS

This work was supported by the National Institute of Mental Health of the National Institutes of Health under the Award Number R01MH111417 and the Dutch Research Council under award number 18969. The content is solely the responsibility of the authors and does not necessarily represent the official views of the National Institutes of Health.

## AUTHOR CONTRIBUTION STATEMENT

Conceptualization: MGBY, WS, AAB, JCWS, NP.

3D VAMOS computational pipeline – including MR signals: MGBY.

BOLD fMRI experiments: WS, AAB, ECAR.

BOLD fMRI data analysis: WS, AAB, ECAR.

Figures design: MGBY.

Writing - original draft: MGBY.

Writing – review and editing: All authors.

Funding acquisition: MvO and NP.

## DISCLOSURE/CONFLICT OF INTEREST

The authors declare that they have no known competing financial interests, conflict of interest or personal relationships that could have appeared to influence the work reported in this paper.

## CODE/DATA AVAILABILITY STATEMENT

The code and data underlying the findings of this study are available from the corresponding author upon request. Access is subject to a nonexclusive, revocable, non-transferable, and limited right to use solely for research and evaluation purposes, excluding any commercial use.

## ABBREVIATIONS

3D: three-dimensional
7T: 7 tesla
BOLD: blood oxygenation level-dependent
CBF: cerebral blood flow
CBV: cerebral blood volume
CMRO2: oxygen metabolism
CSF: cerebrospinal fluid
fMRI: functional magnetic resonance imaging
GE: gradient echo
GM: grey matter
HcT: hematocrit
SE: spin echo
TE: echo time
VAMOS: vascular model based on statistics
WM: white matter

## Notes

### Competing Interest Statement

The authors have declared no competing interest.

